## supplementary material for "Oropouche virus cases identified in Ecuador using an optimised rRT-PCR informed by metagenomic sequencing"

rRT-PCR assay optimisation and validation

Primer (table S1, reverse primer used was Ec2 R) concentrations were tested in multiple combinations at 1 µM, 3 µM, 9 µM and 18 µM. The optimal concentration for both the forward and reverse primer was 18 µM. Probe concentration was tested from 5 µM to 25 µM, with the optimal concentration being 12.5 µM. Magnesium sulphate ($MgSO_4$) concentration was optimised by adding additional $MgSO_4$ to the reaction mix, from none to a maximum of 85 mM. The optimal condition was no added $MgSO_4$. Cross-reactivity to 23 virus species (table S2) and a panel of negative human sera was assessed, no cross-reactions were observed.

| Oligo name | Sequence (5' - 3') | Start position | End position | Length (bp) | Tm (°C) | GC content (%) | Reference |
|---|---|---|---|---|---|---|---|
| OROV F | CATTTGAAGCTAGATACGGACAA | 118 | 140 | 23 | 59 | 39 | Weidmann *et al.* 2003 |
| OROV R | CCATGGGCCTCGATG | 225 | 211 | 15 | 52 | 67 | Weidmann *et al.* 2003 |
| Ec R | CCATGGGCCGCGACG | 225 | 211 | 15 | 57 | 80 | This study |
| Ec2 R | CATCTTTGGCCTTCTTTTRG | 198 | 179 | 20 | 54-56 | 40-45 | This study |
| OROV P | CAATGCTGGTGTTGTTAGAGTCTTCTTCCT | 146 | 175 | 30 | 69 | 43 | Weidmann *et al.* 2003 |

**Table S1.** Oligonucleotides used in the development of the OROV rRT-PCR.

| Virus family | Virus genus | Virus species | Virus acronym |
|---|---|---|---|
| Arenaviridae | Mammarenavirus | Tamiami mammarenavirus | TAMV |
| Flaviviridae | Flavivirus | Powassan virus | POWV |
| Flaviviridae | Flavivirus | West Nile virus | WNV |
| Flaviviridae | Flavivirus | Yellow fever virus | YFV |
| Flaviviridae | Flavivirus | Karshi virus | KSIV |
| Flaviviridae | Flavivirus | Usutu virus | USUV |
| Flaviviridae | Flavivirus | Dengue virus serotype 1 | DENV-1 |
| Flaviviridae | Flavivirus | Dengue virus serotype 2 | DENV-2 |
| Flaviviridae | Flavivirus | Dengue virus serotype 3 | DENV-3 |
| Flaviviridae | Flavivirus | Dengue virus serotype 4 | DENV-4 |
| Flaviviridae | Flavivirus | Zika virus | ZIKV |
| Nairoviridae | Orthonairovirus | Crimean-Congo hemorrhagic fever orthonairovirus | CCHFV |
| Nairoviridae | Orthonairovirus | Issyk-Kul virus | ISKV |
| Peribunyaviridae | Orthobunyavirus | Batai orthobunyavirus | BATV |
| Peribunyaviridae | Orthobunyavirus | La Crosse orthobunyavirus | LACV |
| Peribunyaviridae | Orthobunyavirus | Inkoo virus | INKV |
| Peribunyaviridae | Orthobunyavirus | Tahyna virus | TAHV |
| Phenuiviridae | Phlebovirus | Bhanja virus | BHAV |
| Phenuiviridae | Phlebovirus | Severe fever with thrombocytopenia syndrome virus | SFTSV |
| Phenuiviridae | Phlebovirus | Rift Valley fever phlebovirus | RVFV |
| Togaviridae | Alphavirus | Chikungunya virus | CHIKV |
| Togaviridae | Alphavirus | Mayaro virus | MAYV |
| Togaviridae | Alphavirus | O'nyong-nyong virus | ONNV |

**Table S2.** Viruses included in the rRT-PCR exclusivity testing.

| **Number of mismatches** | **Forward primer** | **Reverse primer (OROV R)** | **Reverse primer (Ec2 R)** | **Probe** |
|---|---|---|---|---|
| **1** | 17 | 20 | 41 | 21 |
| **2** | 2 | 6 | 0 | 0 |
| **>2** | 0 | 0 | 0 | 0 |

**Table S3.** Mismatches to primer sequences seen in an alignment of 149 OROV N gene sequences. Values are the number of sequences with mismatches to the primer/probe sequence.

| Genome position | Genome segment | Gene | Position within gene | D-057 base | D-087 base | D-155 base | D-171 base | D-206 base | D-210 base |
|---|---|---|---|---|---|---|---|---|---|
| 329 | S | N | 285 | G | G | G | G | A | G |
| 551 | S | N | 507 | T | C | C | C | C | C |
| 689 | S | N | 645 | A | G | G | G | G | G |
| 1501 | M | M | 518 | A | A | T | A | A | A |
| 1751 | M | M | 768 | T | T | T | T | C | T |
| 2038 | M | M | 1055 | C | C | C | C | T | C |
| 2230 | M | M | 1247 | A | A | A | A | A | G |
| 2363 | M | M | 1380 | T | C | T | T | T | T |
| 2403 | M | M | 1420 | A | G | G | G | G | G |
| 2810 | M | M | 1827 | G | R* | G | G | G | G |
| 2859 | M | M | 1876 | A | A | A | A | A | G |
| 3290 | M | M | 2307 | T | T | C | T | T | T |
| 4028 | M | M | 3045 | C | C | C | C | C | A |
| 4124 | M | M | 3141 | G | G | A | G | G | G |
| 4313 | M | M | 3330 | T | T | T | T | C | T |
| 4340 | M | M | 3357 | G | A | A | A | A | A |

| | | | | | | | | | |
|---|---|---|---|---|---|---|---|---|---|
| 4490 | M | M | 3507 | T | C | C | C | C | C |
| 6129 | L | L | 747 | T | C | T | T | T | T |
| 6174 | L | L | 792 | A | A | A | A | G | A |
| 6200 | L | L | 818 | A | A | A | A | A | G |
| 6579 | L | L | 1197 | G | G | G | G | A | G |
| 7599 | L | L | 2217 | T | C | C | C | C | C |
| 8865 | L | L | 3483 | G | A | A | A | A | A |
| 9235 | L | L | 3853 | T | T | T | T | T | C |
| 9336 | L | L | 3954 | G | G | A | G | G | G |
| 9571 | L | L | 4189 | A | A | G | A | A | A |
| 9591 | L | L | 4209 | G | A | G | G | G | G |
| 10039 | L | L | 4657 | A | G | G | G | G | G |
| 10737 | L | L | 5355 | A | A | A | G | A | A |
| 10791 | L | L | 5409 | C | C | C | C | C | T |
| 11133 | L | L | 5751 | C | T | C | C | C | C |
| 11208 | L | L | 5826 | C | C | C | C | A | C |
| 11733 | L | L | 6351 | C | C | C | C | T | C |

**Table S4.** SNPs identified between the Ecuadorian OROV genomes. Variant base is shaded grey. * R position = 79% T, 21% C.

| Protein | Isolate | Codon | Consensus AA | SNP AA | R group change |
|---|---|---|---|---|---|
| NSs | D-206 | 88 | C | Y | None |
| M (Gn) | D-155 | 173 | Q | L | Polar / non-polar |
| M (NSm) | D-206 | 352 | A | V | None |
| M (NSm) | D-210 | 416 | K | R | None |
| M (NSm) | D-057 | 464 | A | T | Non-polar / polar |
| M (Gc) | D-210 | 626 | T | A | Polar / non-polar |
| L | D-210 | 273 | D | G | Acidic / non-polar |
| L | D-155 | 1397 | T | A | Polar / non-polar |

**Table S5.** Amino acid variation between Ecuadorian OROV genomes. AA = amino acid. Gn = glycoprotein Gn. NSm = non-structural protein NSm. Gc = glycoprotein Gc. Bunyavirus Gn, NSm and Gc protein positions are taken from GenPept entry AGH07923.1. R group = reactive group.

| OROV strain | S | M | L | Total |
|---|---|---|---|---|
| D-057 | 1 | 0 | 0 | 1 |
| D-087 | 1 | 6 | 5 | 12 |
| D-155 | 2 | 6 | 7 | 15 |
| D-171 | 1 | 2 | 3 | 6 |
| D-206 | nd | 2 | 7 | 9 |
| D-210 | 0 | 2 | 5 | 7 |

**Table S6.** The number of SNPs present in each genome segment (S, M and L), between patient and cultured genome sequences. n/a = no patient genome data available.

| Sample ID | Sex (M/F) | Age | Days of Fever | OROV rRT-PCR (Cq) | Estimated genome copies/ml plasma |
|---|---|---|---|---|---|
| D-057 | M | 35 | 3 | 25.66 | $1.26 \times 10^9$ |
| D-087 | M | 41 | 7 | 36.26 | $9.62 \times 10^3$ |
| D-155 | M | nd | 2 | 25.75 | $1.19 \times 10^9$ |
| D-171 | F | nd | 2 | 26.75 | $6.04 \times 10^8$ |
| D-206 | M | nd | 4 | 30.87 | $1.97 \times 10^7$ |
| D-210 | M | nd | 3 | 29.18 | $9.27 \times 10^7$ |

**Table S7.** OROV positive sample metadata and rRT-PCR results. nd = no data.

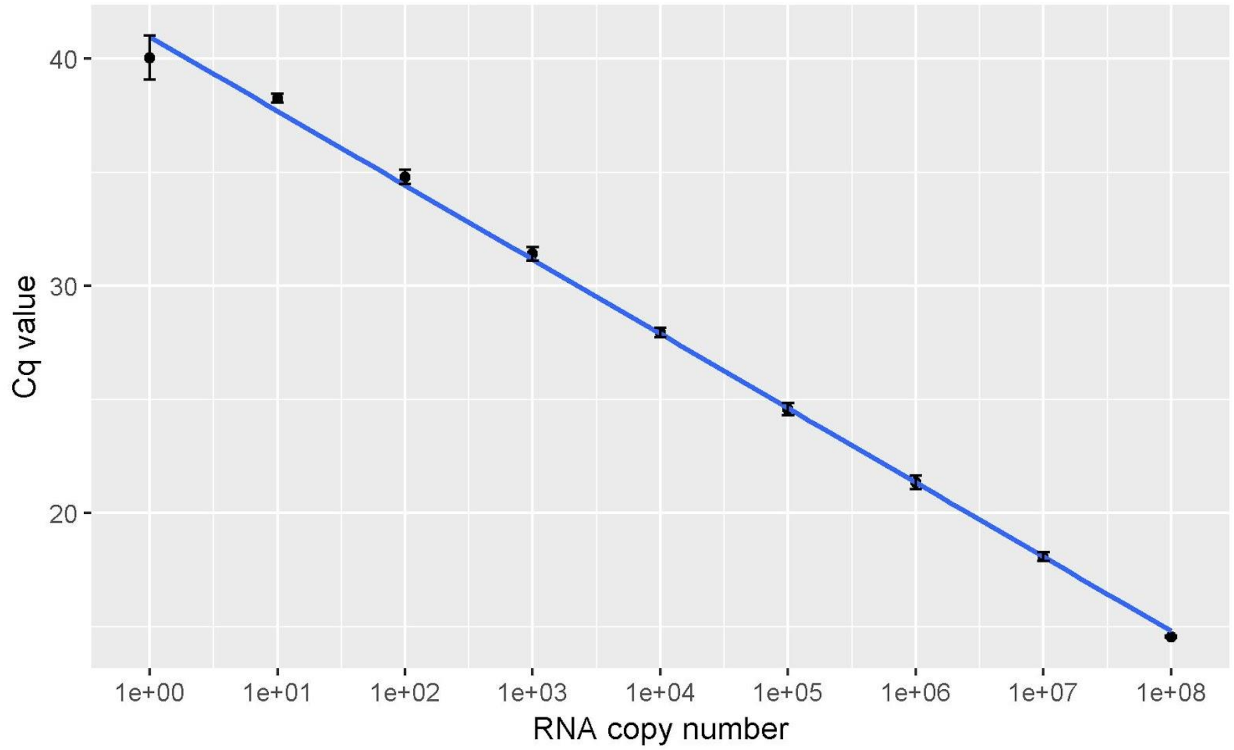


**Figure S1.** Cq value vs OROV RNA copy number, tested in triplicate. Error bars indicate standard deviation. $R^2$ correlation coefficient = 0.9978.

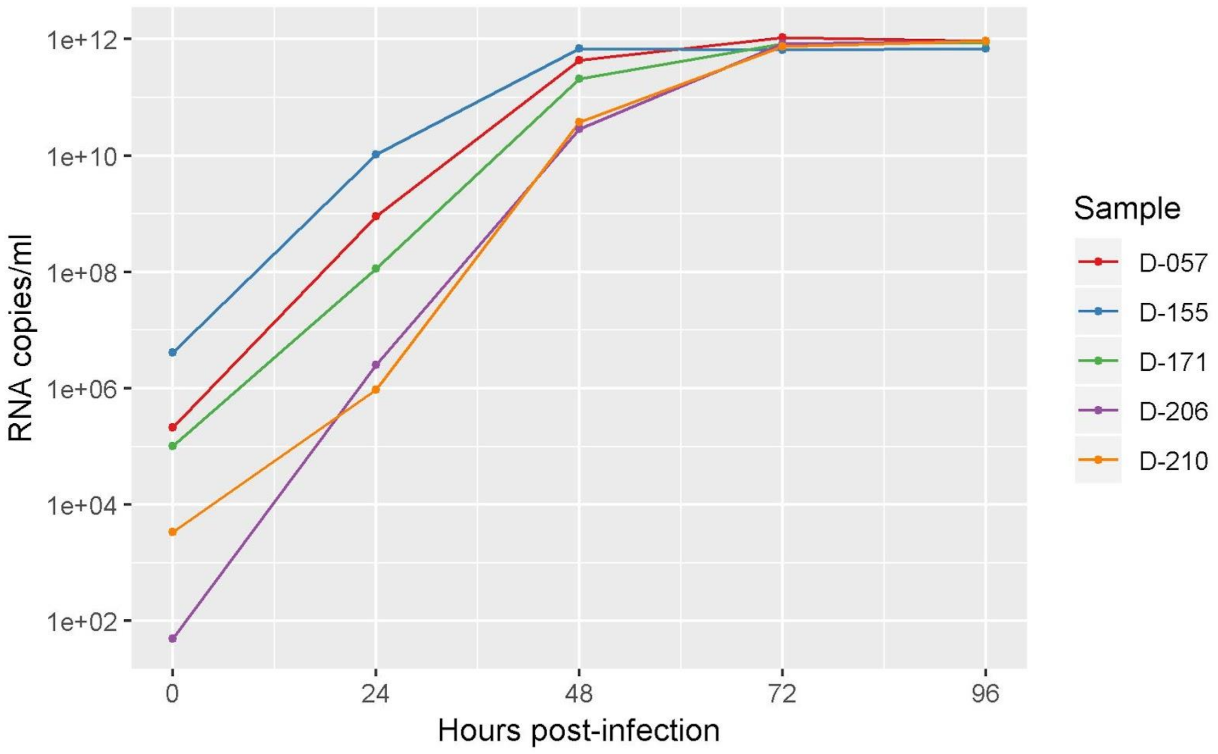


**Figure S2.** Viral genome replication within OROV cultures, isolated from patient plasma in Vero cells.

### Multiplex tiling PCR primer details

Primer details are available as a .csv file at the following link:

Supplementary material. OROV multiplex tiling PCR primers
